# Rab7 activation ameliorates impaired lysosome dynamics and mitophagic flux during chronic iron overload

**DOI:** 10.64898/2026.11.10.765721

**Authors:** Mélanie Mansat, Hye Kyoung Sung, Eddie Tam, Shiraz Anwar, Maria R. Narciso, Gary Sweeney, Roberto J. Botelho

## Abstract

Iron homeostasis is essential for cellular function, yet iron overload is linked to pathologies including cardiovascular disease and metabolic syndrome. Although excess iron has been associated with impaired autophagy and mitochondrial dysfunction, the effects of iron overload on lysosomal dynamics and mitophagy remain poorly defined. In this study, we demonstrate that iron overload disrupts mitochondrial morphology and lysosomal homeostasis, accompanied by iron accumulation in both organelles. While mitophagy is triggered upon iron overload, mitophagic flux is impaired, suggesting a defect in mitophagosome maturation and incomplete clearance. Notably, lysosomes in iron-loaded cells were hyper-acidic and degradative, but became enlarged, perinuclear, and less motile, suggesting that altered lysosomal dynamics may contribute to defective mitophagic clearance. Interestingly, pharmacological activation of Rab7 restores lysosomal motility and mitophagic flux under iron overload conditions. It also improves mitochondrial function and cell viability. Together, these findings indicate that iron overload disrupts organelle homeostasis by impairing lysosomal and mitochondrial dynamics, ultimately arresting mitophagic flux due to altered lysosome dynamics and promoting cell death. Targeted activation of Rab7 mitigates these iron-induced effects, providing a potential therapeutic target for iron overload-induced cell death and tissue damage.

## Introduction

Iron is essential for biological processes like oxygen transport, mitochondrial respiration, and DNA synthesis. For example, in skeletal muscle, iron is a crucial component of myoglobin and mitochondrial oxidative phosphorylation enzymes like Complex I, which respectively stores oxygen and produces ATP (Andrews and Schmidt, 2007; Barakat et al., 2025; Rineau et al., 2021). However, iron homeostasis needs to be tightly regulated because iron excess can lead to extensive cellular damage by generating reactive oxygen species (ROS) through the Fenton reaction. Consequently, this can then cause DNA, lipid, and protein damage, and mitochondrial depolarization, which ultimately cause ferroptosis (Dixon et al., 2012; Hou et al., 2016; Rizzollo et al., 2021). From a pathophysiological perspective, elevated iron levels are observed in cases of iron overload cardiomyopathy, endocrine disorders, liver failure, diabetes, and in patients receiving frequent blood transfusions (Ying Wang et al., 2022). Although iron cytotoxicity is known, the effect of iron overload on the different organelles and the mechanisms leading to ferroptosis are still unclear.

Iron is transported in the bloodstream by transferrin and is endocytosed into cells via several routes, predominantly via the transferrin receptor (TfR). As the transferrin-TfR complexes are delivered into acidic endosomes, the iron ions are released from transferrin (Dautry-Varsat et al., 1983). Once free, the iron is reduced to ferrous iron by metalloreductases like STEAP3 (Ohgami et al., 2006). Membrane transporters such as DMT1 then move the iron to the cytosol (Tabuchi et al., 2000), where it is either stored within ferritin nanocages or translocated to the mitochondrial matrix to synthesize heme and iron-sulfur clusters (Shaw et al., 2006; Wolff et al., 2018). However, excess iron overwhelms iron storage mechanisms leading to cellular damage (Galy et al., 2024). This raises questions about how cells maintain organelle integrity and function when faced with iron overload.

Damaged organelles are cleared via proteostatic mechanisms such as autophagy (Anding and Baehrecke, 2017). For example, mitochondrial membrane depolarization triggers the canonical mitophagy pathway governed by the PINK1/Parkin cascade (Geisler et al., 2010; Matsuda et al., 2010; Narendra et al., 2010). Depolarization causes an accumulation and stability of PINK1 on the mitochondrial outer membrane, which in turn phosphorylates and stabilizes the E3 ubiquitin ligase Parkin on the mitochondrion (Kane et al., 2014; Koyano et al., 2014). Parkin activation leads to protein ubiquitylation and recruitment of the autophagosome formation machinery *via* the LC3 protein, and the formation of mitophagosomes (Narendra et al., 2008). Mitophagosomes then fuse with lysosomes maturing into acidic and degradative mitolysosomes responsible for recycling the damaged mitochondria. Lysosomes are thus critical to sustain autophagic and mitophagic flux and recycle building blocks. Lysosome function and fusogenicity are impacted by their motility and position within cells (Cabukusta and Neefjes, 2018; Korolchuk et al., 2011; Wijdeven et al., 2016). For example, peripherally distributed lysosomes are less acidic than those found perinuclearly (Johnson et al., 2016). Lysosome fusion, motility, and position are in turn governed by several GTPases, particularly the Rab7 GTPase, which modulates FYCO1 and RILP, which are respective adaptors of kinesin and dynein motors (Baba et al., 2019; Cabukusta and Neefjes, 2018; Langemeyer et al., 2018). Indeed, the Rab7 GTPase is critical for Parkin-mediated mitophagic flux (Jin et al., 2022; Yamano et al., 2018). Interestingly, Rab7 also regulates mitochondria-lysosome contact sites and mitochondria fission (Wong et al., 2018). Overall, lysosomal function is essential for maintaining mitochondria quality and for removing damaged mitochondria.

Previously, we discovered that excess iron triggered autophagy but that over time this was disrupted due to impaired autophagosome-lysosome reformation (Jahng et al., 2019). Mitophagy can be triggered upon iron overload, as evidenced by an accumulation of damaged mitochondria and increased levels of the autophagic LC3-II and p62 (Jee et al., 2026; Tam et al., 2024; Zhang et al., 2020a). Here, we postulated that iron may impair mitophagic flux by disrupting lysosome function. Indeed, we found increased mitophagy and abated mitophagic flux in cells chronically overloaded with iron. Surprisingly, lysosomes were hyperacidic and hyperdegradative; instead, they became more static and redistributed heavily to the perinuclear region of the cell, which likely reduced their fusogenicity. Pharmacological activation of Rab7 increased lysosomal motility, restored mitophagic flux, and improved the overall cell viability.

## Results

### Mitochondria accumulate iron and become fragmented during chronic iron exposure

Iron overload impairs insulin sensitivity, glucose uptake, and autophagy (Jahng et al., 2019; Sung et al., 2023). In addition, a role of mitochondrial iron in cell death was described, but the mechanism remained poorly defined (Jee et al., 2026; Tam et al., 2024; Yuan et al., 2016; Zhang et al., 2020a). To better understand this, we used the previously established iron overload model by treating L6 skeletal cells with 250 µM ferrous labile iron for up to 24 h and investigated the effects on mitochondrial properties and their quality control systems.

Firstly, using MitoFerroFluor (MFF), which is a fluorescent probe that accumulates in mitochondria and is quenched by iron, we showed that its fluorescence was significantly reduced in iron-treated cells relative to control cells, indicating an accumulation of iron in mitochondria (Fig. 1A, B). Secondly, cells loaded with iron displayed mitochondria that gained in sphericity, and became smaller and less networked (Fig. 1C-F), suggesting that iron causes mitochondrial fragmentation. As mitochondria are dynamic organelles that undergo fusion and fission, their fragmentation could be due to a decrease in fusion and/or an increase in fission. To assess which process may be affected by iron overload, we used Western blotting to quantify the levels of fusion-promoting Mitofusins and Opa1, and fission-driving Fis1 and phosphorylated-Drp1. While Mitofusin1 (Mfn1) and p-DRP1 levels were unchanged (Fig. 1G, H, K), iron overload appears to decrease the levels of Mitofusin2 (Mfn2) and Opa1 fusion proteins (Fig.1 G, I-J), while increasing the amount of Fis1 fission proteins relative to resting cells (Fig.1G, L). These changes are consistent with altered mitochondrial dynamics, reflecting enhanced mitochondrial fission and/or reduced fusion rates, in agreement with previous studies (Tam et al., 2024; Yuan et al., 2016). Given the previously observed accumulation of oxidative stress in mitochondria (Tam et al., 2024) and higher fragmentation (here and in (Tam et al., 2024), we postulated that mitophagy may be engaged in iron-loaded cells.

**Figure 1.**
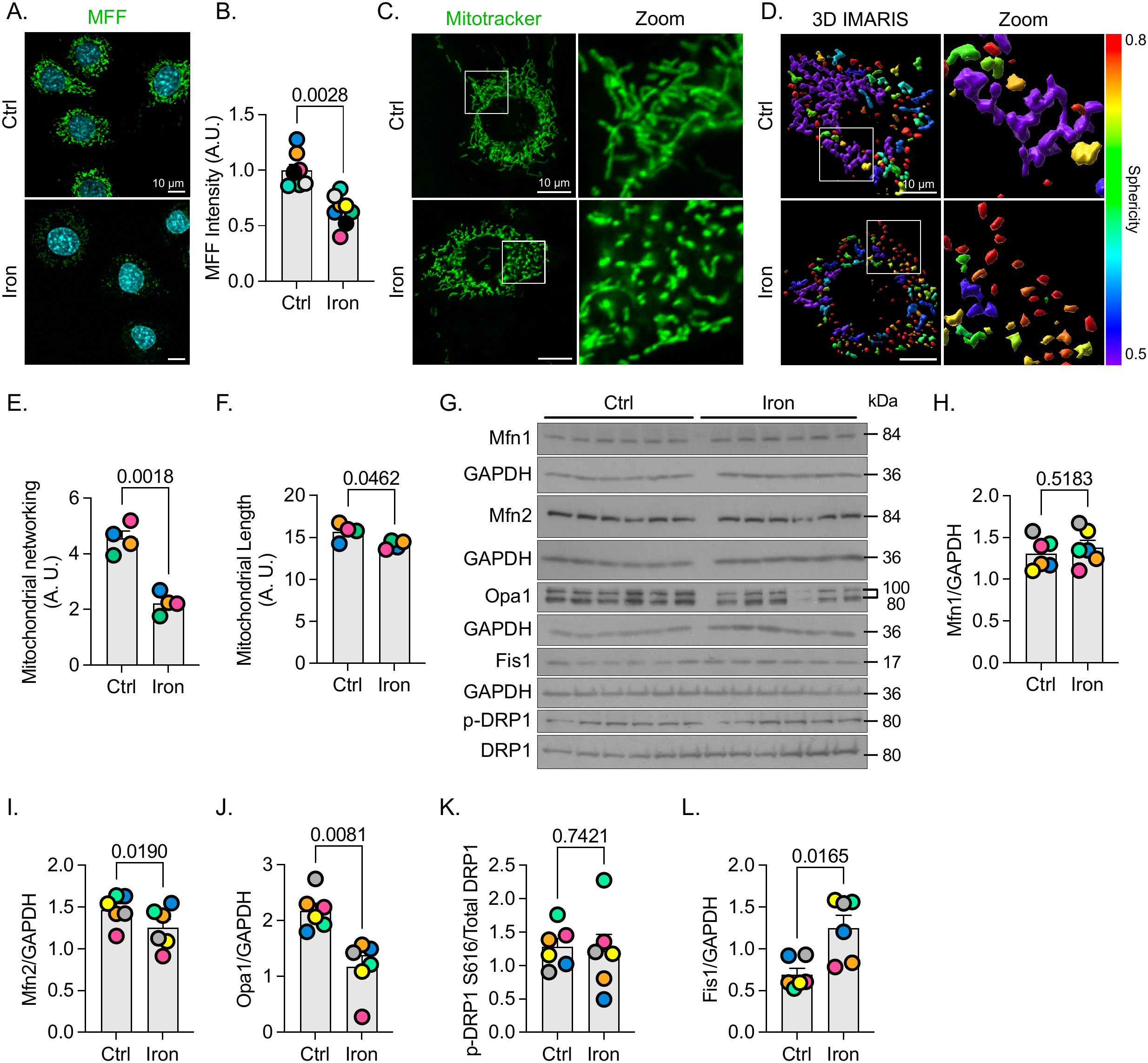
Chronic iron overload disturbs mitochondria morphology. L6 cells were treated with or without iron for 24 h. **A.** Representative confocal microscope images of L6 cells labelled for mitochondrial iron using MitoFerroFluor (MFF, green) and for nuclei with DAPI (cyan). **B.** Relative MFF intensity in control and iron-loaded cells. Data are shown as mean ± SEM from 8 independent experiments, where at least 10 cells per condition per experiment were quantified. Data were tested using paired Student’s t-test. **C.** Representative confocal microscope images of L6 cells labelled for mitochondria using Mitotracker. **D.** 3D reconstruction of Mitotracker-labelled cells and visualized for sphericity of the mitochondria **E, F.** Quantification of the mitochondrial networking (E) and length (F). Data are shown as mean ± SEM from 4 independent experiments, where at least 10 cells per condition per experiment were quantified. Data were tested using paired Student’s t-test. **G-L.** Western blot of the whole cell lysates of L6 cells treated or not with iron for 24 h, each lane represent an independent experiment, (G) and the quantification of Mfn1 (H), Mfn2 (I), Opa1 (J) and Fis1 (L) bands relative to GAPDH and phospho-DRP1 (S616) band relative to DRP1 (K). Data are shown as mean ± SEM from 6 independent experiments and tested using a paired Student’s t-test.

### Chronic iron overload triggers Parkin-dependent mitophagy

We tested if iron overload would trigger Parkin-dependent mitophagy in L6 cells overexpressing mCherry-Parkin. Cells were kept resting, or exposed to FCCP (a known trigger of Parkin-dependent mitophagy), or overloaded with iron. We then quantified the percent of cells with only cytosolic Parkin, mixed cytosolic and punctate distribution, or predominantly Parkin-labelled puncta, as well as the number of Parkin-labelled puncta per cell. While 50% of control cells displayed cytosolic Parkin only, this dropped to <5% in iron-loaded and FCCP-treated cells, and a corresponding augmentation in the other two categories (Fig. 2A, B); similarly, there was a major increase in the number of mCherry-Parkin-labelled puncta per cell (Fig. 2A, C). Additionally, Parkin co-localization with TOMM20 increased in iron and FCCP-treated cells relative to control cells (Fig. 2A, D). Collectively, these data suggest that iron overload triggers Parkin-dependent mitophagy in L6 cells. We previously suggested that cells loaded with iron over 24 h exhibited reduced Parkin on mitochondria isolated with a mitochondria isolation kit (Tam et al., 2024). However, in retrospect, this kit may not have isolated mitochondria enclosed in autophagosomes, thus reducing the apparent amount of Parkin on isolated mitochondria.

**Figure 2.**
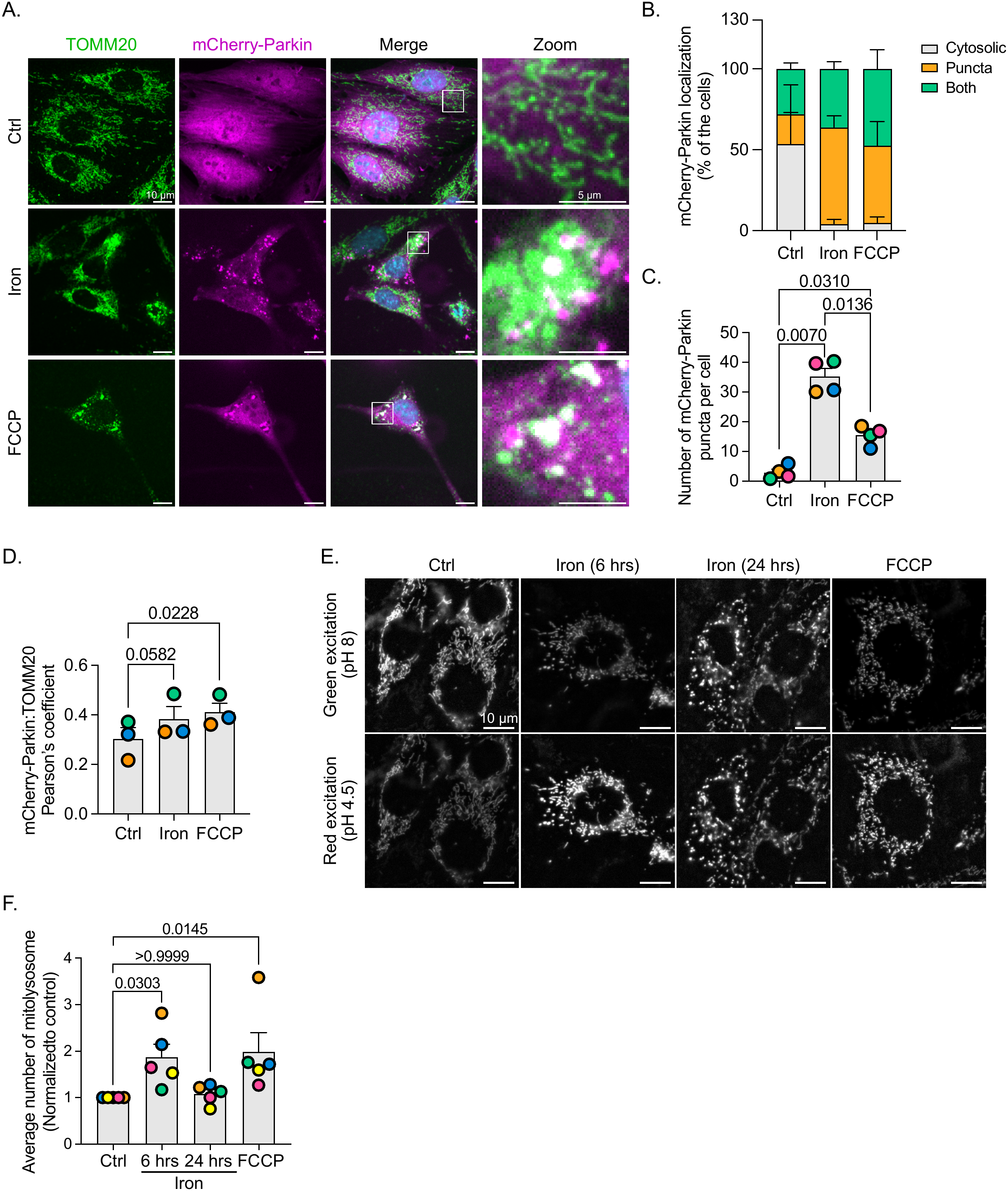
Chronic iron overload triggers mitophagy but blocks mitophagic flux. L6 cells were treated with or without iron for 6 or 24 h, or with FCCP to trigger mitophagy. **A.** Representative confocal micrographs of L6 cells overexpressing mCherry-Parkin (magenta) and labelled for mitochondria using TOMM20 (green) and for the nuclei with DAPI (cyan). **B, C, D.** Quantification of the percentage of cells with cytosolic or mixed mCherry-Parkin distribution, or mostly puncta (B), the number of mCherry-Parkin puncta per cell (C) and the Pearson’s coefficient between mCherry-Parkin and Tomm20 (D). Data are shown as mean ± SEM from 3 independent experiments, where at least 30 cells per condition per experiment were quantified. Data were tested using One-way ANOVA and Tukey’s (C) and Dunnett’s (D) multiple comparisons test. **E.** L6 stable cell line expressing mt-Keima were imaged live to follow the mitophagic flux. Shown is the emission after green (optimally excitation at alkaline pH) or red (optimally excited at acidic pH) excitation. Merge of the two emissions is shown. **F.** Normalized average red signal after subtraction of the green signal as an indicator of mitophagosome acidification. Data are shown as mean ± SEM from 5 independent experiments, where at least 30 cells per condition per experiment were quantified. Data were tested using Friedman test and Dunn’s multiple comparisons test.

### Mitophagic flux is deficient in cells overburdened with iron

After mitophagy, mitophagosomes need to fuse with lysosomes to degrade and recycle the mitochondrial cargo, a process referred to as mitophagic flux (Youle and Narendra, 2011). Fusion with lysosomes acidifies the mitophagosomes, which can be estimated in living cells by expressing the mt-Keima probe (Kelly et al., 2024; Liu et al., 2021; Sun et al., 2015). L6 cells stably expressing an inducible mt-Keima were treated with vehicle, FCCP, or with iron for 6 h (acute treatment) or 24 h (chronic overload). Mitophagic flux was assessed by counting the number of mitolysosomes per cell, as described previously (Kelly et al., 2024).

Cells exposed to iron for 6 h only displayed an increase in red-excitation over green- excitation relative to resting cells and comparable to FCCP-treated cells (Fig. 2E, F), implying mitophagosome acidification and flux. In comparison, cells overloaded with iron for 24 h favoured the green-based excitation relative to 6 h iron and FCCP-treated cells (Fig. 2E, F), indicating failed acidification of mitophagosomes after prolonged iron loading. Collectively, our observations indicate that iron overload triggers Parkin-dependent mitophagy, but these fail to mature. These observations are consistent with previous results showing that chronic iron overload promotes LC3-based autophagy but fails to mature into autolysosomes (Jahng et al., 2019; Sung et al., 2023). We postulated that this may occur due to damage to lysosomes.

### Iron accumulates but does not rupture lysosomes

Free ferrous iron likely enters cells by membrane transporter activity at the plasma membrane and by endocytosis (Gunshin et al., 1997; Hentze et al., 2010). In the latter case, iron then traffics to endosomes where it might transfer to mitochondria (Shaw et al., 2006). However, excess iron may be misdirected and accumulate in and damage lysosomes, which may then obstruct mitophagic flux. Indeed, cells exposed to 24 h of ferrous iron displayed a 5-fold increase in FerroOrange fluorescence, an iron indicator, in Lysotracker-labelled lysosomes (Fig. 3A, B), consistent with previous observations (Fakih et al., 2008). Considering this, we tested for lysosome damage by expressing GFP-galectin-3, a cytosolic protein that rapidly assembles on the exposed luminal membrane of ruptured lysosomes forming puncta (Papadopoulos and Meyer, 2017). Surprisingly, there was no difference in the number of cells labelled with galectin-3-GFP puncta in control and iron-loaded group (Fig. 3C, D). In comparison, ∼80% of cells exhibited galectin3-GFP puncta when treated with LLoMe, a potent lysosome damaging agent – compared to only 20-25% for the other conditions (Fig. 3C, D). Overall, these data suggest that iron does not cause coarse lysosome membrane damage within the conditions used and in L6 cells.

**Figure 3.**
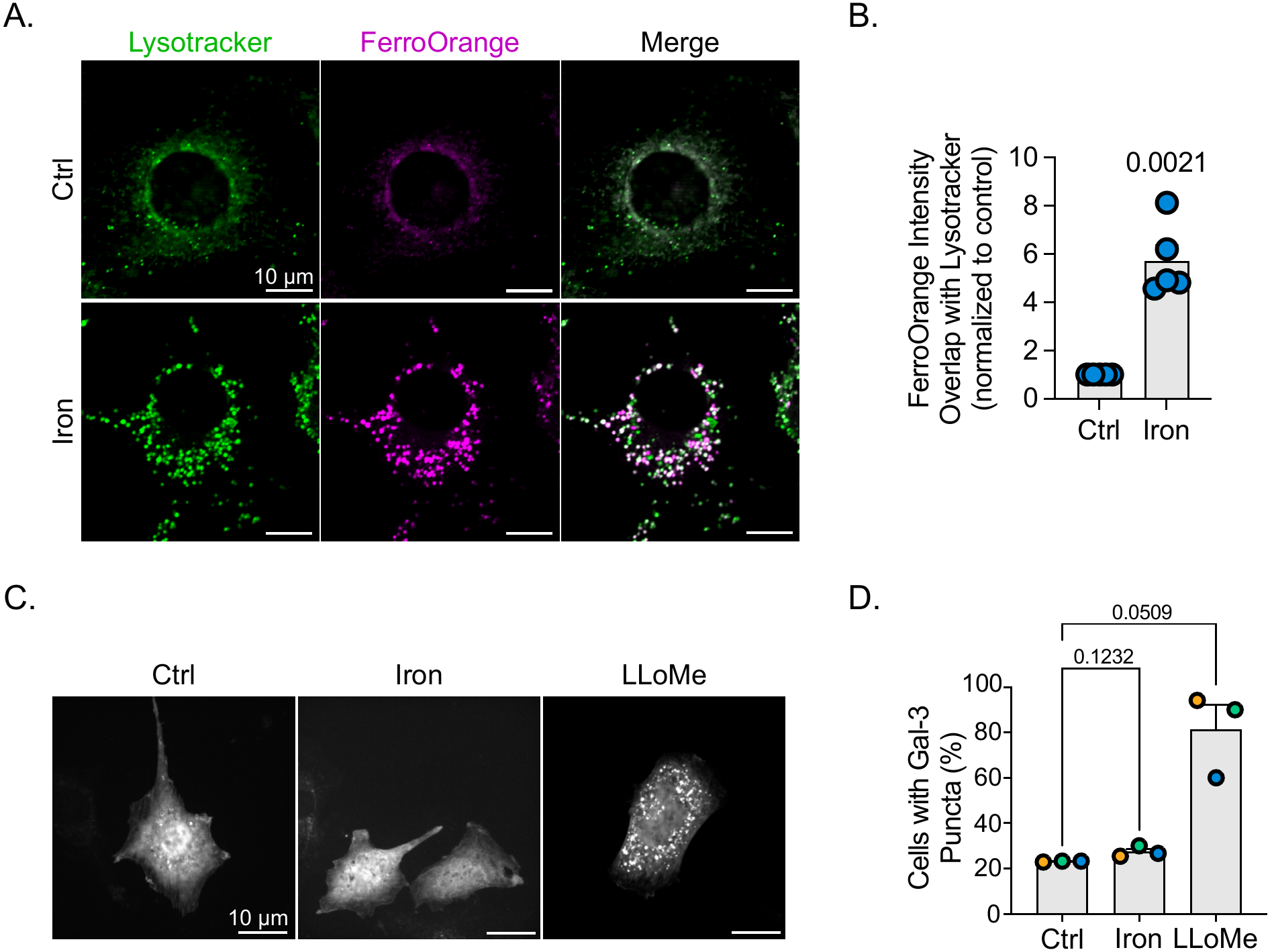
Lysosomal membrane integrity is maintained upon iron overload. **A.** Representative confocal images of resting and iron-loaded L6 cells labelled for lysosomes using Lysotracker (green) and for iron using FerroOrange (magenta). **B.** Quantification of the overlap between FerroOrange labeling and Lysotracker, normalized to control. Data are shown as mean ± SEM from 5 independent experiments, where at least 10 cells per condition per experiment were quantified. Data were tested using a one-sample Student’s t-test. **C.** Representative confocal images of L6 cells overexpressing GFP-Galectin-3 under resting, iron-loaded, or LLoMe-exposed conditions. **D.** Quantifications of the percentage of cells with GFP-Galectin-3 puncta. Data are shown as mean ± SEM from 3 independent experiments, where at least 30 cells per condition per experiment were quantified. Data were tested using One-way ANOVA and Dunnett’s multiple comparisons test.

### Lysosomes are hyperacidic and hyperdegradative in cells overloaded with iron

We next sought to determine if iron overload perturbed lysosomal pH and/or degradative capacity, which might then blunt mitophagic flux. L6 cells were loaded with pHrodo-red-dextran, a pH-sensitive dye used to track the endolysosomal trafficking, challenged with vehicle or iron for 6 or 24 h, and then pHrodo fluorescence was measured. Interestingly, resting cells and those loaded with iron for 6 h had similar pHrodo fluorescence, which was reduced upon incubation with bafilomycin A, a V-ATPase inhibitor (Fig. 4A, B). Surprisingly, cells exposed to iron for 24 h exhibited higher pHrodo fluorescence relative to resting cells, indicating hyperacidification (Fig. 4A, B); consistent with this notion, bafilomycin A completely dampened the hyperfluorescence of pHrodo in cells that consumed iron for 24 h (Fig. 4A, B) showing that this was pH-dependent.

**Figure 4.**
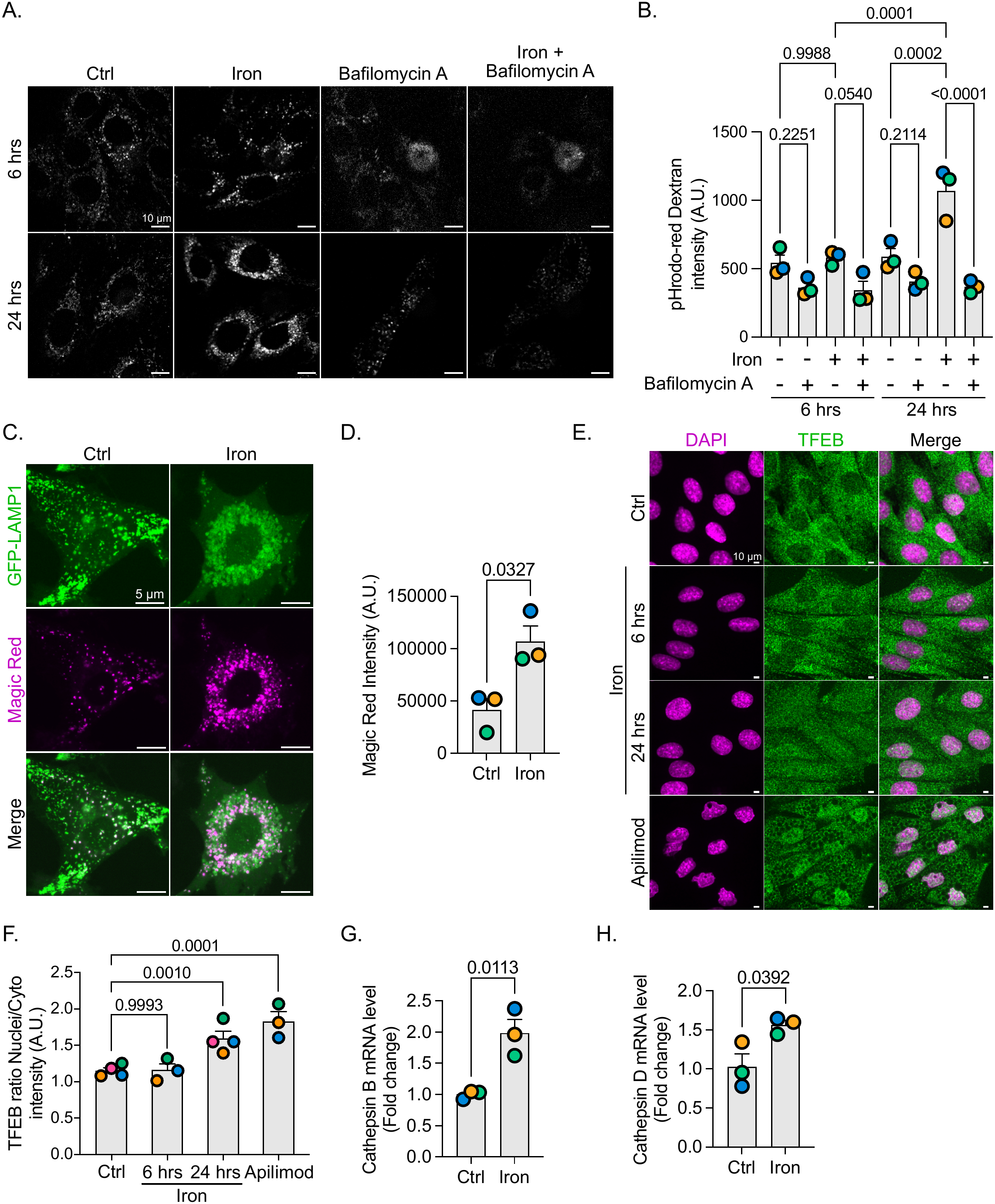
Long term iron exposure hyperactivates lysosome degradation and acidification. **A.** Confocal micrographs of L6 cells pre-loaded with pHrodo red-dextran as an indicator of lysosomal pH and then maintained at rest or loaded with iron for 24 h. Bafilomycin A was used to demonstrate pH-sensitivity of pHrodo red-dextran. **B.** Quantification of the pHrodo red-dextran intensity. Data are shown as mean ± SEM from 3 independent experiments, where at least 40 cells per condition per experiment were quantified. Data were tested using One-way ANOVA test and Šídák’s multiple comparisons test. **C.** Confocal microscope images of resting and iron-loaded L6 cells overexpressing GFP-LAMP1 (green) and labelled using Magic Red (magenta) to follow lysosomal Cathepsin L activity. **D.** Quantifications of the Magic Red intensity. Data is shown as mean ± SEM from 3 independent experiments, where at least 40 cells per condition per experiment were quantified, and tested using paired Student’s t-test. **E.** Images of resting, iron-loaded (6 and 24 h), and apilimod-treated L6 cells labelled for TFEB (green) and the nuclei with DAPI (magenta) to follow TFEB translocation from the cytosol to the nuclei. **F.** Quantification of the fluorescence intensity ratio between nuclear to cytosolic TFEB. Data are shown as mean ± SEM from 3 independent experiments, where at least 60 cells per condition per experiment were quantified. Data were tested using One-way ANOVA test and Dunnett’s multiple comparisons test. **G, H.** Quantification of Cathepsin B (G) and Cathepsin D (H) mRNA level in L6 cells treated or not with iron for 24 h. Data are shown as mean ± SEM from 3 independent experiments tested with an unpaired Student’s t-test.

We then evaluated proteolytic activity of lysosomes by staining resting and iron-loaded cells with Magic Red, a marker of Cathepsin L activity. As with pHrodo fluorescence, cells chronically overloaded with iron presented superior Magic Red intensity compared to control cells (Fig. 4C, D), suggesting hyper-degradative lysosomes. We anticipated that hyper-acidification and hyper-degradation could partly be due to activation of TFEB and enhanced expression of lysosomal enzymes triggered by iron-based stress. Indeed, relative to resting and short iron exposure (6 h), there was an enrichment in nuclear TFEB in cells incubated with iron for 24 h that was comparable to apilimod, a PIKfyve inhibitor known to promote TFEB (Fig. 4E, F; (Choy et al., 2018; Gayle et al., 2017)). Consistent with TFEB activation, there was a corresponding increase in Cathepsin B and D mRNA levels (Fig. 4 G, H). Overall, these data intimate that lysosomes are hyper-acidic and hyper-degradative in L6 cells chronically loaded with iron.

### Iron-loaded lysosomes are swollen, perinuclear, and less motile

Our data till now suggest that lysosomes are not grossly damaged and in fact are more acidic and degradative than their control counterparts. However, as we performed these experiments, we suspected that lysosome dynamics were altered. We thus sought to quantify morphology, distribution, and motility in iron-loaded cells. First, lysosomes were visibly enlarged in iron-loaded cells relative to resting lysosomes (Fig. 4C). Second, using shell-analysis, lysosomes redistributed to the perinuclear region in iron-loaded cells (Fig. 4C, 5A, 5B). Third, there was a marked decrease in lysosome track displacement and total distance travelled in iron-loaded cells relative to vehicle group (Fig. 5B-D) and as depicted by kymographs (Fig. 5E). This suggests that lysosome dynamics are defective in cells chronically exposed to iron, which may impair fusion with target organelles like mitophagosomes and autophagosomes.

**Figure 5.**
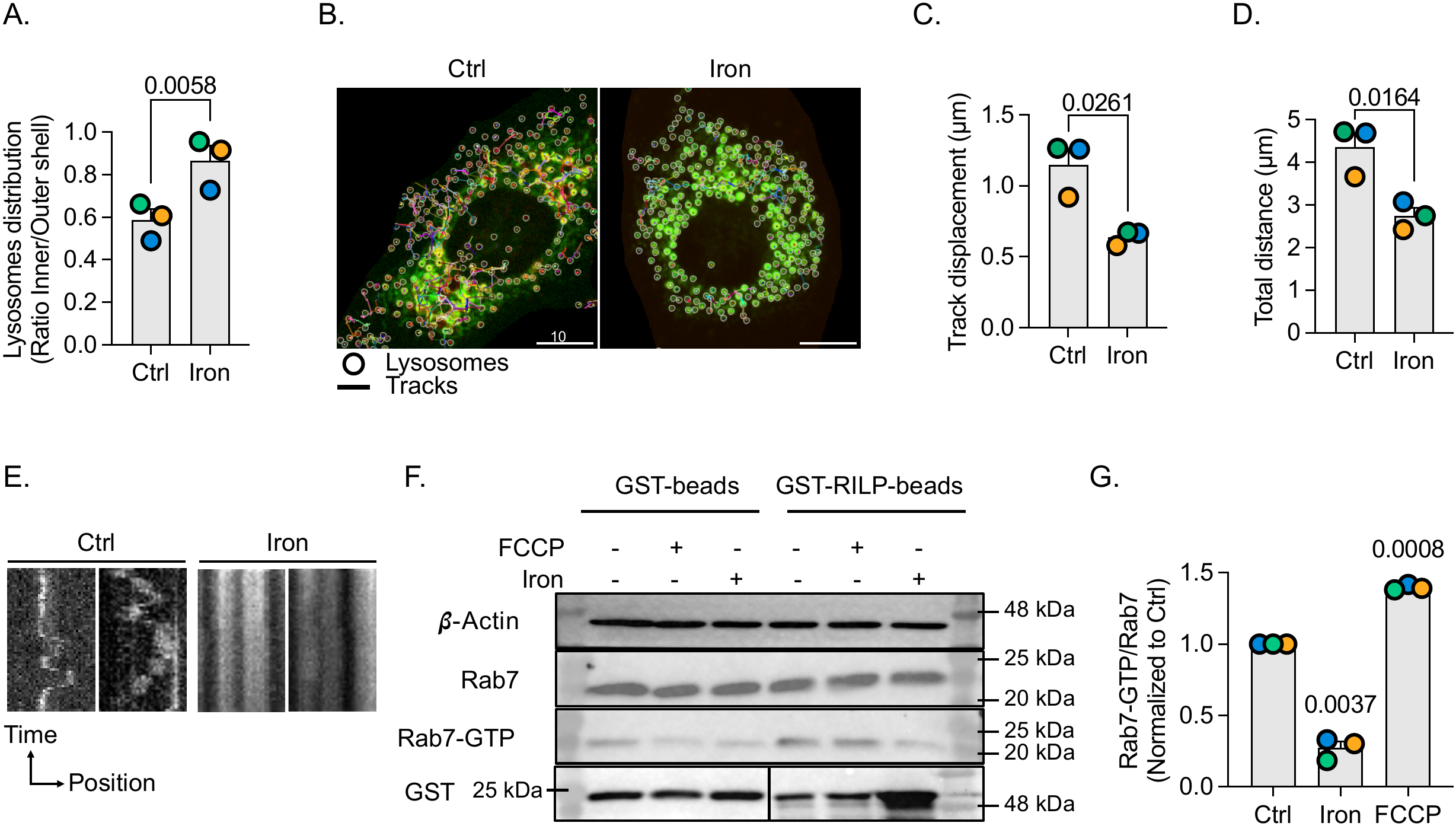
Long term iron exposure alters lysosome distribution and motility. Lysosome enlargement and perinuclear distribution of lysosomes in L6 cells loaded with iron relative to resting cells. **A.** Lysosome distribution quantification as the ratio between perinuclear lysosomes (inner shell) to peripheral lysosomes (outer shell). **B.** Representative image of the lysosomal track displacement overlay. “Circles” denote individual lysosomes, and “lines” are the different tracks followed by each lysosome over a 2-minute period, with one image every 2 seconds. **C, D.** Quantification of the lysosomal track displacement (C) and total distance (D) over a 2-minute period, with one image every 2 seconds. For A, C, D, data are shown as mean ± SEM from 3 independent experiments and tested by a paired Student’s t-test. **E.** Representative kymographs of lysosomal movement in resting and iron-loaded cells. **F.** Western blotting of affinity precipitated GTP-bound Rab7 (Rab7-GTP) after GST-RILPC33 beads incubation of L6 cell lysates treated with vehicle, iron, or FCCP. GST and GST-RILPC33 levels are shown (GST). Whole cell inputs are shown in the top two Westerns (β-actin and total Rab7). **G.** Quantification of the ratio between Rab7-GTP over total Rab7. Data are shown as mean ± SEM from 3 independent experiments tested with one sample t-test against a hypothetical value of 1.

Given these changes to lysosome dynamics, we suspected that iron may alter the activity of master lysosome regulators like the Rab7 GTPase, which recruits motor proteins and fusion factors (Jin et al., 2022; Wong et al., 2018; Yamano et al., 2018). To test this, we quantified GTP-Rab7 affinity precipitation using beads loaded with the GST-RILPC33 chimera (Margiotta et al., 2017). As FCCP was also reported to increase Rab7 activity, we used it as a positive control. Compared to resting cells, there was a significant drop in GTP-Rab7 recovered when cells were loaded with iron for 24 h (Fig. 5F, G). Overall, iron overload significantly disrupts lysosome dynamics without disturbing membrane integrity, alkalinizing the pH, and hindering hydrolase activity.

### Pharmacological activation of Rab7 rescues iron-induced changes to lysosome motility and distribution

Whilst other regulators of lysosomes may be disturbed, we postulated that the reduced Rab7 activation may contribute to altered motility and/or abated fusion in iron-loaded cells, impairing mitophagic flux. To test this, we turned to ML098, a reported pharmacological activator of Rab7 (Jin et al., 2022; Surviladze et al., 2010a). Indeed, ML098 exposure increased the levels of Rab7 by immunofluorescence relative to vehicle cells (Sup. Fig. S1A, B). We then examined the effects on lysosome dynamics in iron-loaded cells exposed to ML098. As before, iron overburden blunted lysosome motility (Fig. 6A-C, column 1 vs. column 3). Interestingly, we did not observe an effect on lysosome dynamics when cells were treated with ML098 alone (Fig. 6A-C, column 1 vs. column 2). Nonetheless, cells co-treated with iron and ML098 were rescued for the lysosome motility defects incurred by iron overload as measured by lysosome track displacement and total distance travelled (Fig. 6A-C). Moreover, while iron-alone caused perinuclear lysosomal redistribution, this was reversed when ML098 was added to cells, becoming comparable to the resting cell counterparts (Fig. 6D). Intriguingly, ML098 could not rescue all lysosomal changes forced by iron overburden; for example, iron-loaded L6 cells retained higher Magic Red fluorescence with or without ML098 (Fig. 6E, F).

**Figure 6.**
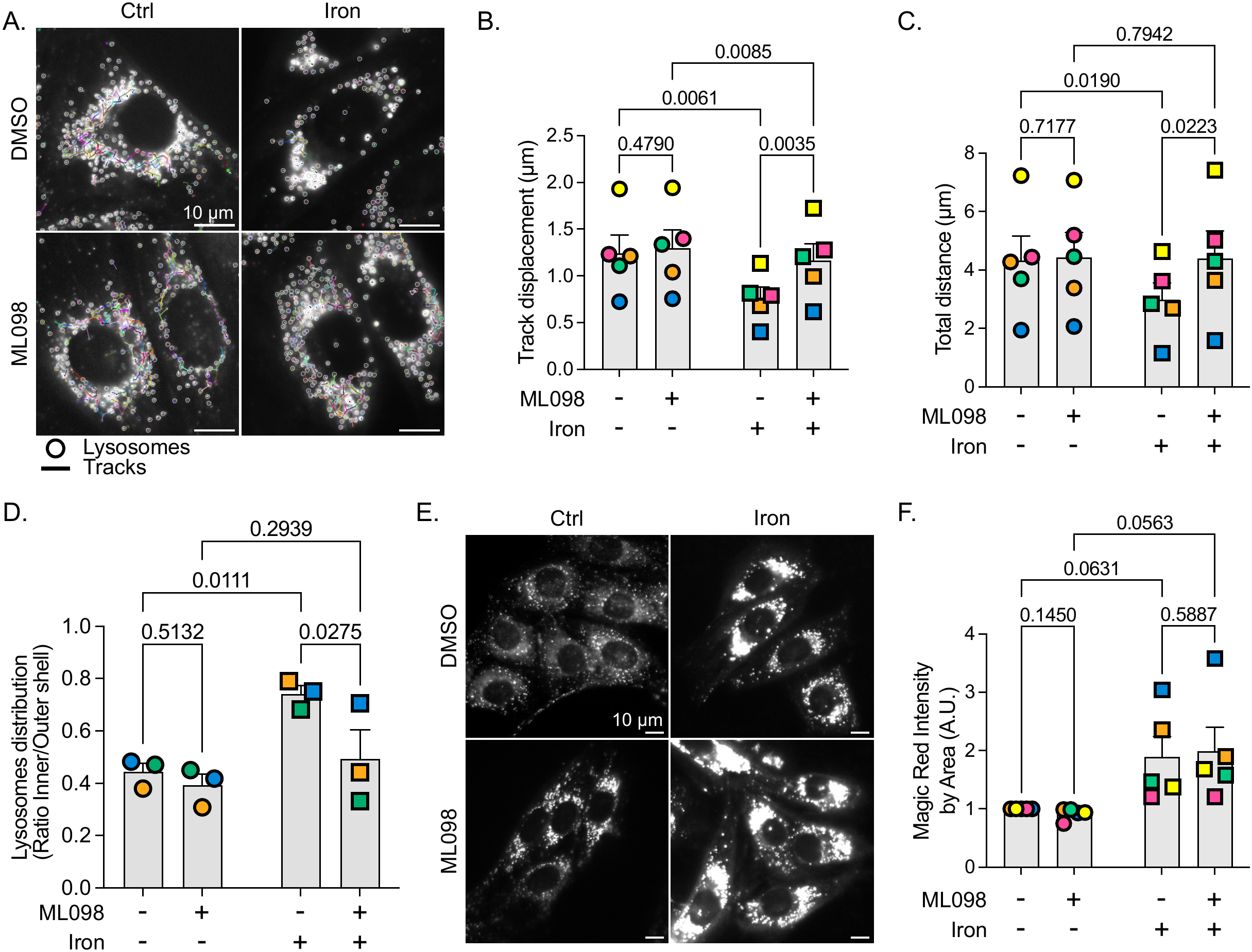
Pharmacological activation of Rab7 improves lysosomal motility in iron-loaded cells. **A.** Representative image of the lysosomal track displacement overlay. “Circles” denote individual lysosomes, and “lines” are the different tracks followed by each lysosome over a 2-minute period, with one image every 2 seconds. **B, C.** Quantification of the lysosomal track displacement (B) and total distance (C) over a 2-minute period, with one image every 2 seconds. Data is shown as mean ± SEM from 5 independent experiments where at least 60 cells per condition per experiment were quantified. Data were tested using Two-ways ANOVA and Fisher’s LSD. **D.** Quantifications of the ratio between the lysosome in the inner shell (around the nuclei area) and the outer shell (away from the nuclei area). Data is shown as mean ± SEM from 3 independent experiments, where at least 15 cells per condition per experiment were quantified. Data were tested using Two-ways ANOVA and Fisher’s LSD. **E, F.** Representative confocal microscope images of L6 cells labelled with Magic Red to follow lysosomal Cathepsin L activity (E) and quantifications of the Magic Red intensity (F). Data is shown as mean ± SEM from 5 independent experiments where at least 60 cells per condition per experiment were quantified. Data were tested using Two-ways ANOVA and Fisher’s LSD.

### Pharmacological activation of Rab7 rescues iron-impaired mitophagic flux

Given the improved motility and redistribution of lysosomes induced by ML098, we then tested if ML098 could revert the mitophagic flux defect in iron-stressed cells. First, ML098 alone had no apparent effect on mt-Keima fluorescence properties (Fig. 7A, B). As before, iron-loaded cells displayed mt-Keima fluorescence consistent with neutral pH, with mitochondria appearing more fragmented (Fig. 7A, B). In comparison, there was a marked recovery of the mt-Keima acidic excitation in iron-loaded cells co-exposed with ML098 relative to iron-loaded cells alone (Fig. 7A, B). This suggests that ML098 promotes mitophagic flux in cells overloaded with iron. Supporting these data, while both LC3-II and p62 expression were significantly increased in iron-loaded cells compared to the control, these were reduced if ML098 was present (Fig. 7C-E). VDAC, an outer mitochondrial membrane protein, was used as a mitochondrial loading control to assess changes in mitochondrial protein abundance. Taken together, these data suggest that pharmacologically induced Rab7 activation promotes lysosome motility and mitophagic flux in cells chronically overloaded with iron, which may aid in mitochondrial recycling and recovery.

**Figure 7.**
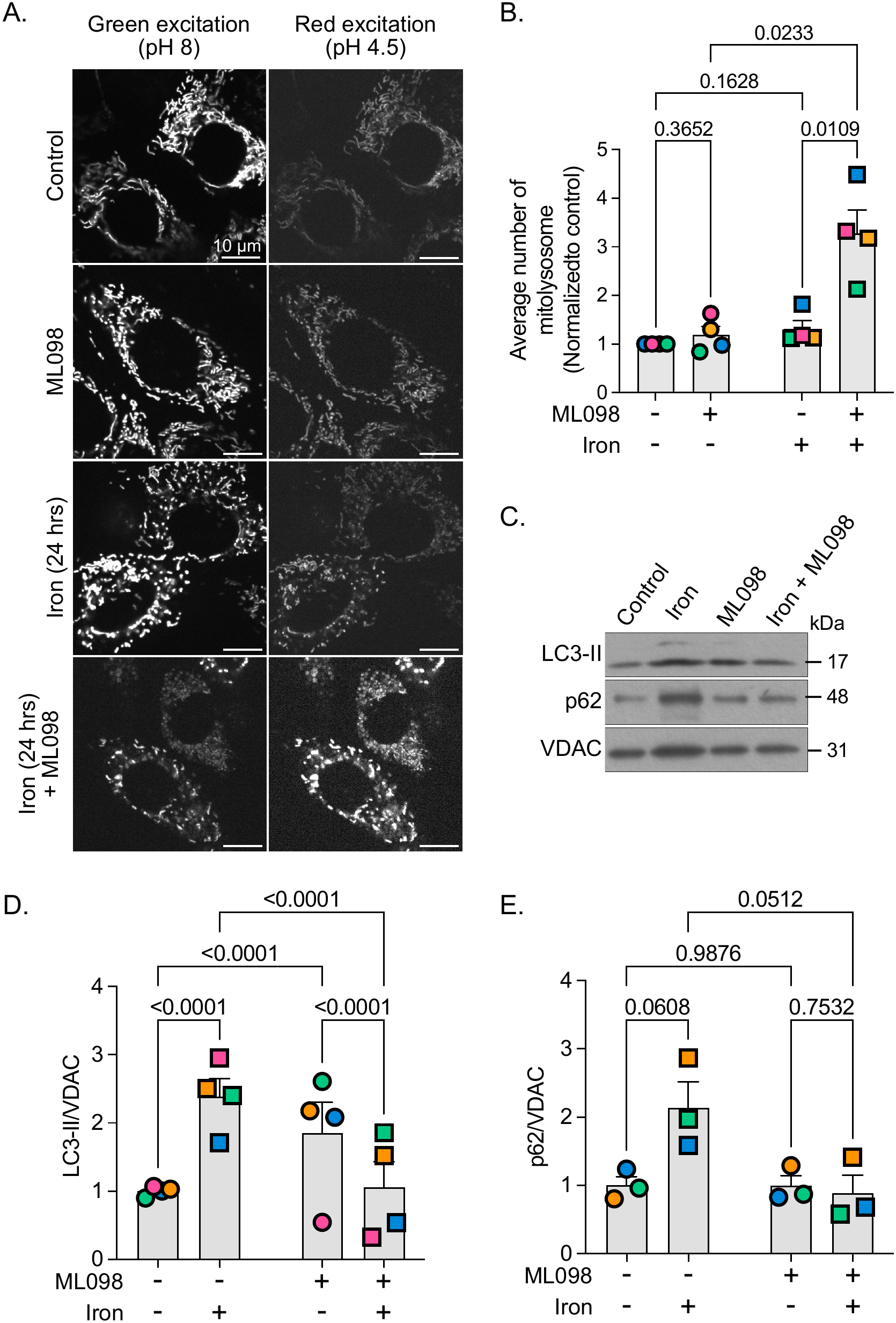
Rab7 activation restores mitophagic flux. **A.** Micrographs of L6 cells stably expressing mt-Keima in resting and iron-overloaded conditions with and without 100 nM ML098. Images are emission at 600/50 nm after excitation with 470 nm (approximate optimal for alkaline excitation) and 555 nm (approximate optimal for acidic milieu excitation) **B.** Normalized average mitolysosome area per cell as described in methods. Data are shown as mean ± SEM from 4 independent experiments, where at least 30 cells per condition per experiment were quantified. **C-E.** Western blot of the whole cell lysates of L6 cells treated with iron for 24 h and/or with ML098 for indicated proteins (C) and the quantification of LC3-II (D) and p62 (E) bands relative to VDAC. For B, D and E, data are shown as mean ± SEM from 3-4 independent experiments and tested using two-way ANOVA and Fisher’s LSD.

### ML098 mitigates the toxic effects of chronic iron overload in cells

We postulated that the enforced Rab7 activation and corresponding boost in mitophagic flux might improve mitochondrial quality and cell survival in cells overstrained with iron. To test this, we assessed two key indicators of mitochondrial quality: mitochondrial membrane potential and lipid peroxidation. Excess iron can promote lipid peroxidation and compromise mitochondrial membrane integrity, contributing to the loss of mitochondrial membrane potential. Therefore, we assessed mitochondrial membrane depolarization using TMRE, a fluorescent dye that accumulates in polarized mitochondria. Iron overload significantly decreased TMRE mean intensity relative to control cells (Fig. 8A, B). Importantly, ML098 rescued the iron-induced drop in TMRE labelling to levels comparable to control cells, though ML098 on its own had minimal effects (Fig. 8A, B). To determine if mitochondrial lipid peroxidation was rescued by ML098, L6 cells were labelled using MitoCLox, a mitochondrial lipid peroxidation sensor. Excess iron potently increased the MitoCLox fluorescence ratio compared to control cells, demonstrating lipid peroxidation (Fig. 8C, D). Remarkably, ML098 decreased MitoCLox labelling when co-administered with iron (Fig. 8C, D), showing that Rab7 activity mitigated the accumulation of mitochondrial damage.

**Figure 8.**
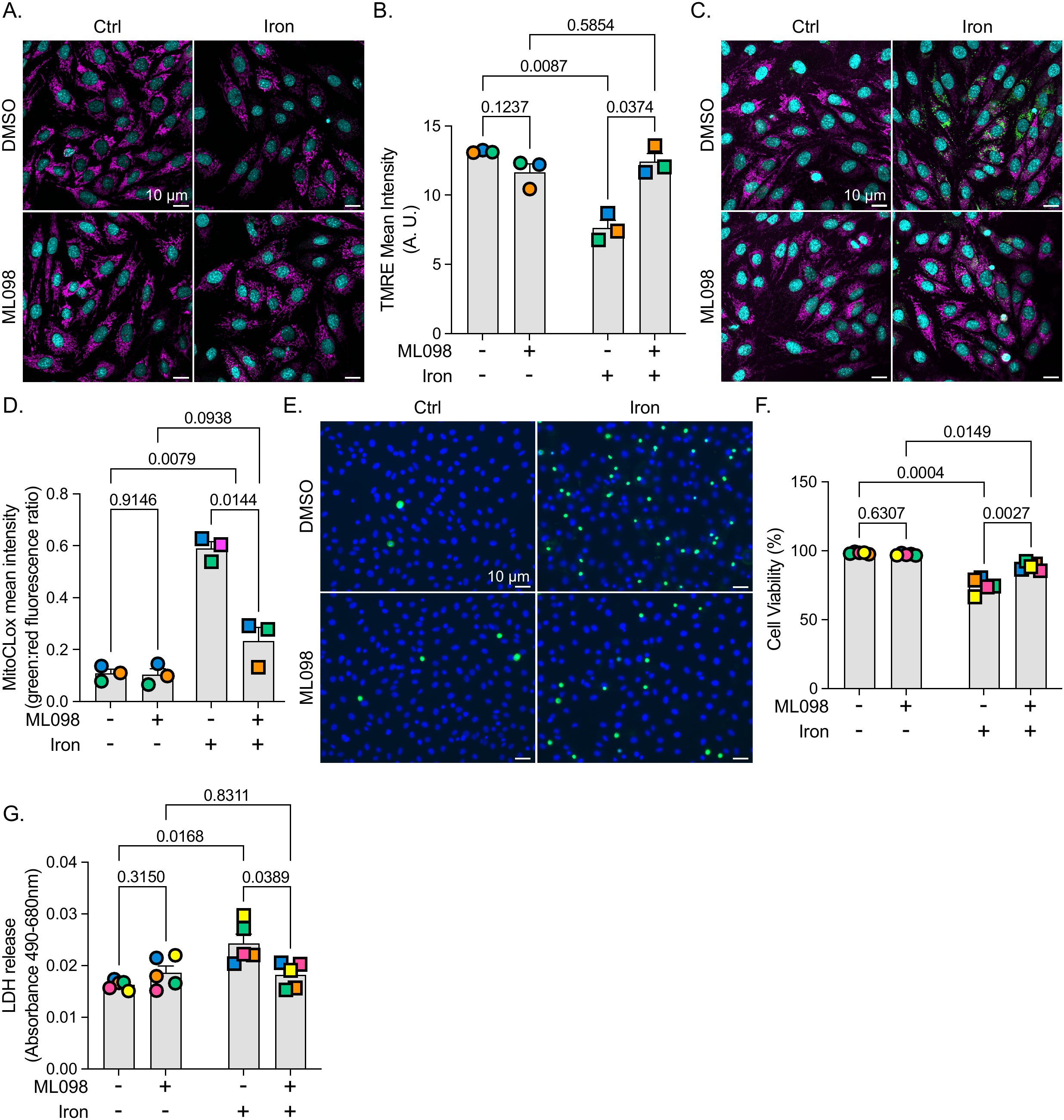
Rab7 activation improves chronic iron overload cell survival. L6 cells were treated with or without iron for 24 h, and/or with 100 nM ML098 to induce Rab7 activation. **A.** Representative confocal microscope images of L6 cells labelled using TMRE dye (magenta) to measure mitochondrial membrane potential and for nuclei using Hoechst (cyan). **B.** Quantification of the TMRE intensity per cell. **C.** Representative confocal microscope images of L6 cells labelled using MitoCLox dye (magenta, green) to measure mitochondrial lipid peroxidation and for nuclei using Hoechst (cyan). **D.** Quantification of the MitoCLox intensity ratio (green:red). **E.** Representative confocal microscope images of L6 cells labelled using ReadyProbe Cell viability kit containing Hoechst (nuclei, blue) and propidium iodide (dying cells, green). **F.** Quantification of percent cell viability. **G.** Quantification of LDH released by L6 cells after the indicated treatments. For B, D, F, and G: Data are shown as mean ± SEM from 3 independent experiments (B, D) or five experiments (F, G) and analysed by two-way ANOVA and Fisher’s LSD.

Lastly, the degree of cell death was assessed in L6 cells by measuring the fluorescence of the Readyprobe cell viability imaging kit which distinguishes viable from non-viable cells, and by quantifying lactate dehydrogenase (LDH) release into the extracellular medium. Upon iron treatment, cell viability was significantly decreased to ∼75% and LDH release was significantly increased compared to control conditions (Fig. 8E-G). In comparison, iron-loaded cells treated with ML098 showed improved cell viability (∼85%) and a significant decrease in LDH release compared to iron-alone (Fig. 8E-G). Overall, these results suggest that ML098-mediated Rab7 activation can mitigate iron overloading damage to mitochondria and cell function, possibly by boosting mitophagosome turnover that is arrested in iron overloaded cells.

## Discussion

The mechanisms underlying the damage and progression to cell death caused by chronic iron overload are incompletely understood (Cho et al., 2026; Zhao et al., 2024). Chronic iron overload promotes autophagy (Fernández et al., 2016a; Jahng et al., 2019; Sung et al., 2023; Yunyang Wang et al., 2022; Zhou et al., 2021), but thwarts autophagic flux (Fernández et al., 2016b; Jahng et al., 2019). Similarly, iron overload has been well documented to alter mitochondrial dynamics and function in various models, leading to oxidative stress and cell death (Gao et al., 2010; Park et al., 2015; Tam et al., 2024, 2022; Wang et al., 2024; Zhao et al., 2024). Subsequently, we showed that overexpression of mitoNEET, a mitochondrial transmembrane protein involved in iron–sulfur cluster biology, protected cells against iron-induced stress and ferroptosis (Tam et al., 2024). Here, we expanded on these observations by showing that mitochondrial damage triggers Parkin-dependent mitophagy. However, we propose that mitophagic flux is obstructed due to impaired lysosome motility and fusogenicity, rather than ruptured lysosomes or a defect in lysosomal acidification and degradative capacity. Supporting this model, we showed that activation of the Rab7 GTPase with ML098 improved lysosomal motility, promoted mitophagic flux, attenuated the accumulation of iron-induced mitochondrial damage, and reduced cellular death. This establishes Rab7 and related lysosome-based pathways as potential therapeutic pathways to combat iron-induced tissue damage.

### Iron overload effects on mitochondria quality and mitophagy

Mitochondria are a major destination of labile iron, which can enter cells via membrane transporters like DMT1 and by endocytosis of transferrin (Dautry-Varsat et al., 1983; Tabuchi et al., 2000; Wolff et al., 2018; Zhao et al., 2024). Labile iron is then buffered in mitochondria via mitochondrial ferritin and frataxin (Campanella et al., 2009; Corsi et al., 2002; Hara et al., 2020; O’Neill et al., 2005; Zhao et al., 2024). However, excess iron overwhelms the iron buffering system of mitochondria, which triggers the Fenton reaction to generate excess ROS that damage mitochondrial DNA, proteins and lipids; this compromises mitochondrial functions including their membrane potential and fission-fusion dynamics (Dixon et al., 2012; Gao et al., 2010; Tam et al., 2024, 2024, 2022; Zhao et al., 2024). These deleterious effects of iron overload on mitochondria have been observed in variety of cellular and physiological models including cardio-vascular, hepatic, lung, pancreatic, and skeletal models (Bacon and Britton, 1989; Gao et al., 2010; Jee et al., 2026; Masini et al., 1989; Tam et al., 2024; Wang et al., 2024; Zhang et al., 2020a), and in iron overload pathologies such as hereditary hemochromatosis, β-thalassemia, Friedreich’s ataxia, and cancer (Cho et al., 2026; Jee et al., 2026; Thakerngpol et al., 1996; Zhao et al., 2024). Hence, our observations on iron-induced alteration of mitochondrial dynamics and function in skeletal cells (Figs. 1, 7, and 8) are thus entirely consistent with the current picture of the disruptive impact of iron excess on mitochondria. Consequently, mitochondrial quality control mechanisms are engaged during iron excess, of which mitophagy is an important one.

Here, we provide evidence for potent iron-induced Parkin-mediated mitophagy (Fig. 2), which is typically triggered by mitochondrial depolarization that sequentially stimulates the kinase PINK1 and the E3 ligase Parkin (Matsuda et al., 2010; Narendra et al., 2010). Given the oxidative damage and loss of membrane potential in iron-loaded cells (Fig. 8), we anticipate that this is the primary mechanism by which iron overload stimulates Parkin-labelling of mitochondria in L6 cells. In fact, there are multiple links between conditions that accumulate iron or induce ferroptosis and mitophagy response (Chang et al., 2024; Chen et al., 2023; Li et al., 2025, 2026; Tam et al., 2024; Wang et al., 2025). However, iron overload was also proposed to reduce mitophagy in some models contributing to cell death (Jee et al., 2026; Zhang et al., 2020b). This may indicate a non-universal response to iron overload, or differences in the models of iron overload or mitophagy, or a discrepancy in interpreting between mitophagy initiation, completion, and/or turnover. For example, in our previous work, we suggested that iron overload induces mitophagy, but we observed reduced Parkin-levels associated with isolated mitochondria using a fractionation kit – in retrospect, that interpretation may be confounded by the changes to mitochondria enclosed partially or completely within mitophagosomes that may not have co-fractionated with intact mitochondria leading to an apparent drop in Parkin levels in isolated mitochondria in iron-loaded cells (Tam et al., 2024).

Post mitophagy, mitophagosomes fuse with lysosomes to mature, degrade, and recycle cargo and membranes – this is mitophagic flux, which is functional within the first 6 h of iron overload. However, our data using mt-Keima and p62/LC3II levels suggests a blockade in mitophagosome maturation/flux under chronic iron overload (Figs. 2, 7). This is consistent with autophagic flux arrest observed in iron-burdened cells (Fernández et al., 2016a; Fu et al., 2023; Jahng et al., 2019; Kao et al., 2016). We had previously attributed this autophagic flux defect to an arrest in autophagic lysosome reformation since most lysosomes were labelled with LC3 (Jahng et al., 2019). However, as described below, additional lysosome changes occur that may also thwart mitophagic flux.

### Iron overload effects on the lysosomal system

Given the defective mitophagic flux, we postulated that iron accumulated in lysosomes causing damaged and rupture. Indeed, iron overload was previously suggested to rupture lysosomes in macrophages (Das et al., 2026; Li et al., 1998), hepatocytes (Myers et al., 1991), and in epithelial cells (Zhang et al., 2023). However, while iron accumulates in lysosomes, we did not observe lysosomal rupture in L6 skeletal cells as detected by GFP-galectin-3 (Fig. 3); instead, lysosomes were more acidic and degradative (Fig. 4). This hyper-state of lysosomes may be partly driven by activation of TFEB, a transcription factor that scales up lysosomal and autophagic activity under stresses such as starvation and infection (Sardiello et al., 2009; Settembre et al., 2013; Visvikis et al., 2014).

While lysosomes appeared hyper-functional for acidification and degradation, they were nevertheless swollen, perinuclear, and static in iron-loaded cells (Figs. 4, 5). We previously observed some of these changes; however, we had also noted that LAMP1^+^-structures were labelled with LC3, leading us to propose that autophagy was triggered, but unresolved due to autophagic lysosome reformation defect (Jahng et al., 2019). However, an emerging alternative is that iron damages lysosomes without rupture, triggering *Conjugation of Atg8 to Single membranes* (CASM); in CASM, mild lysosome damage is known to trigger lipidation and association of LC3 onto lysosomal membranes (Corkery and Wu, 2026; Haukaas et al., 2026; Reid et al., 2022); but as with autophagic and mitophagic flux, this may fail in iron-loaded cells accumulating altered lysosomes that are static and non-fusogenic. Nevertheless, we must note here that others have proposed deacidification of lysosomes in iron-loaded cells, along with swelling and increased LAMP2 expression (Fernández et al., 2016b) – these differences may depend on the cell and iron-loading model, or approaches to measure lysosome properties.

Regardless, the observed changes to lysosome properties we make here may be due to reduced GTP-loaded Rab7 in iron-loaded cells (Fig. 5). While iron excess may affect other factors like the Arl8b GTPase, the fact that ML098, an allosteric activator of Rab7 (Surviladze et al., 2010b), promotes lysosome motility, rescues mitophagic flux, improves mitochondria quality, and aids in cellular survival, suggests that Rab7 inactivation is a key defect in iron overloaded cells (Figs. 6, 7, an 8). Rab7 activation with ML098 was also shown to boost mitophagy and protect cells during ischemic heart failure (Sun et al., 2025), diabetic cardiomyopathy (Ke et al., 2024), pulmonary hypertension (Piper et al., 2024), and promoted autophagic flux in osteonecrosis (Dong et al., 2023). Nevertheless, this may not be a universal mechanism since Fernandez *et al.,* by expressing Rab7 constitutively active and dominant negative mutants, and using an inhibitor of Rab7, suggest that Rab7 activity promotes TPCN channels to induce cell death in iron-loaded cells (Fernández et al., 2016b). We currently do not know the source of this discrepancy.

Lastly, several mechanisms may account for the altered Rab7 activity during iron overload. One possibility involves the regulation of mitochondria-lysosome contact sites (Wong et al., 2018). Rab7 is implicated in the formation and regulation of these contacts, which contribute to the coordination of mitochondrial and lysosomal functions, and mitophagy (Heo et al., 2018; Jin et al., 2022). The mitochondrial Rab7 GTPase-activating protein TBC1D15, which interacts with Fis1 at the mitochondrial surface, participates in the regulation of mitochondria-lysosome contacts and mitochondrial dynamics (Yamano et al., 2018; Yu et al., 2020). The reduced activation of Rab7 upon iron overload may be due to changes in the dynamics of mitochondria-lysosomes contacts (Yamano et al., 2018; Yu et al., 2020).

## Conclusion and limitations

Overall, we propose that iron overload in skeletal cells stresses and damages two organellar systems: lysosomes and mitochondria. Lysosomes do not rupture, but instead are hyper-acidic and hyper-degradative, swollen and static. Additionally, mitochondria are also damaged and marked for degradation by mitophagy. However, mitophagosomes fail to undergo flux due to static lysosomes. As a result, cells enter a program of cell death due to futile clearance of damaged organelles. Kickstarting lysosome motility by activating Rab7 pharmacologically appears to help clear damaged organelles to extend the lifespan of cells. While this study provides mechanistic insights into iron overload-induced lysosomal dysfunction and mitophagic arrest, several limitations warrant consideration. First, and a challenge in the field, are the differences related to cell and iron-loaded models that present discrepancies, whether grounded on physiological and technical distinctions. Additionally, while we present evidence for the PINK1/Parkin axis of mitophagy, other forms of mitophagy may be relevant during iron overload (e.g., BNIP3; (da Silva Rosa et al., 2021)). Lastly, while we propose that ML098 promotes Rab7 activity, the actual effects may be indirect by modulating other factors like Arl8b GTPase and contact sites (Kumar et al., 2022; Wong et al., 2018).

## Materials and Methods

### Cell culture, transfection, and plasmids

The L6 male rat myoblast cell line (ATCC CRL-1458; American Type Culture Collection, Manassas, VA, USA) was cultured at 37°C and 5% CO_2_ in AMEM supplemented with 10% heat-inactivated fetal bovine serum (FBS; Wisent Inc., Saint-Jean-Baptiste, QC, Canada), which we refer to as growth medium. L6 cells were passaged every 2-3 days by lifting them using 0.1% trypsin, resuspended in growth medium, and seeded at 1.5 million cells in a T75 flask; cells were discarded after 20 passages. For experiments, cells were either seeded in 4-well chambered polymer coverslips and grown to 60% confluency, or in 6-well plates and grown to 90% confluency, or at 100,000 cells per well in a 12-well plate and treated the next day in the treatment medium (AMEM supplemented with 0.5% heat-inactivated FBS). If the cells needed to be transfected, they were seeded at 50,000 cells per well in 12-well plate and then treated with Lipofectamine 2000 transfection reagent (Invitrogen, Waltham, MA, USA) following manufacturer instructions. Briefly, for a cell medium volume of 1 mL, 2.4 µg of plasmid was mixed with 1.2 µL of Lipofectamine 2000 in 100 µL of OptiMEM for 15 min before being added to the cells’ medium. Cells were washed the next morning, and subsequent treatments were started the day after. Plasmids used in this work are: pEGFP-LAMP1-SBP (a gift from Juan Bonifacino (Guardia et al., 2019), Addgene plasmid #120172); mCherry-Parkin (a gift from Richard Youle (Narendra et al., 2008), Addgene plasmid #23956); pEGFP-hGal3 (a gift from Tamotsu Yoshimori (Maejima et al., 2013), Addgene plasmid #73080).

### Generation of non-clonal L6 cells stably expressing mt-Keima (L6::mt-Keima cells)

The mitochondrial targeted Keima fluorescent protein (mt-Keima) coding sequence was derived from the mKeima-Red-Mito-7 (a gift from Michael Davidson; Addgene plasmid #56018) by in-frame fusion of the COX8A mitochondrial signal peptide sequence to mKeima. The insert was synthesized by GenScript using SfiI restriction sites and cloned into pSBtet-Hyg, a Sleeping Beauty transposon vector containing an inducible gene-of-interest cloning cassette and hygromycin selection marker (a gift from Eric Kowarz (Kowarz et al., 2015), Addgene plasmid #60508). L6 cells were co-transfected as described above in a 6-well plate using 3 µg of pSBtet-Hyg-mtKeima and 0.3 µg of pCMV(CAT)T7-SB100 (a gift from Zsuzsanna Izsvak (Mátés et al., 2009), Addgene plasmid #34879) plasmids. Cells were washed 24 h after and selected the next day using 200 µg/mL hygromycin B (Gibco, Waltham, MA, USA) for 3 days. Once the cells reached 60% confluency, they were passaged in a T-25 and then in a T-75 flask before freezing and storing in liquid nitrogen. L6::mt-Keima cells were propagated and manipulated as the original L6 parental cells above but with 100 µg/mL hygromycin.

### Iron overload and pharmacological treatments

L6 cells were treated with 250 µM iron (FeSO_4_-7H_2_O, BioShop, Burlington, ON, Canada) for 6 h or 24 h in treatment medium. To induce Rab7 activation, cells were treated with 100 nM ML098 (Cayman Chemicals, Ann Arbor, MI, USA) for 30 min before the iron treatment. For some live-experiments, 200 nM bafilomycin A (Cell Signaling Technologies, Danvers, MA, USA) was used 1 h prior to imaging. To induce mitophagy, cells were treated for 6 h with 10 µM FCCP (Abcam, Cambridge, UK). To induce TFEB nuclear translocation, 100 nM apilimod (Selleck Chemicals, Houston, TX, USA) was used for 4 h. To induce lysosomal membrane rupture, cells were treated with 0.5 mM L-Leucyl-L-Leucine methyl ester (LLoMe, Cayman Chemicals) for 1 h.

### Fluorescent labelling with iron probes

To follow mitochondrial ferrous iron (Fe^2+^) accumulation via fluorescence quenching, MitoFerroFluor (MFF; kindly provided by Dr. John J. Lemasters) was utilized as previously detailed (Tam et al., 2022). At the experimental endpoint, cells were washed three times with Phosphate-Buffered Saline (PBS) and incubated with 2 µM MFF and 100 nM Hoechst for 30 min in PBS at 37 °C, protected from light. Following three subsequent PBS washes, live cells were imaged using a Nikon A1 confocal microscope fitted with an incubation chamber (37 °C, 5% CO_2_). The mean fluorescence intensity of MFF was quantified using ImageJ software.

To follow iron accumulation in lysosomes, cells were simultaneously incubated with 50 nM LysoTracker Deep Red (Thermo Fisher Scientific, Waltham, MA, USA), 100 nM MitoTracker Green (Thermo Fisher Scientific), 1 µM FerroOrange (Dojindo Molecular Technologies, Gaithersburg, MD, USA), and a nuclear stain (Thermo Fisher Scientific) for 30 min in serum-free AMEM at 37 °C, protected from light. After three washes, cells were imaged live via a Nikon A1 confocal microscope under environmental control (37 °C, 5% CO_2_). Manders’ overlap/correlation coefficients were determined using Nikon NIS-Elements Analysis software (Melville, NY, USA).

### Fluorescent labelling with mitochondrial probes

For mitochondria morphology, cells were washed three times and incubated with 100 nM MitoTracker Green (Thermo Fisher Scientific) and the HCS Blue NuclearMask stain (Thermo Fisher Scientific) for 30 min in serum-free AMEM at 37 °C, protected from light. After three additional washes, live-cell imaging was performed using a Nikon A1 confocal microscope equipped with an environmental incubation chamber (37 °C, 5% CO_2_).

For mitochondrial membrane potential measurements, cells were incubated with 50 nM TMRE (Thermo Fisher Scientific) and the HCS Blue NuclearMask stain for 30 min at 37 °C, protected from light. Cells were washed three times prior to immediate live-cell confocal imaging. For mitochondrial lipid peroxidation analysis, cells were loaded with 100 nM MitoCLox (Lumiprobe, Westminster, Maryland, USA) and a nuclear stain for 30 min at 37 °C, protected from light. According to the manufacturer’s instructions, the dye-containing media was removed and replaced with fresh, dye-free medium without washing, followed immediately by live-cell imaging.

To follow mitolysosome formation, L6::mt-Keima cells were seeded in a 12-well plate in glass coverslips and mt-Keima expression was immediately induced using 500 ng/mL of doxycycline (Sigma-Aldrich, Burlington, MA, USA). The live-cell mt-Keima signal was obtained with a Quorum spinning-disk confocal using two filters: 470 nm excitation “green” for neutral-pH, and 555 nm excitation “red” for acidic-pH.

### Fluorescent labelling with lysosomal probes

To follow lysosomal pH, L6 cells were challenged with pHrodo-Red-Dextran 10,000 MW (Invitrogen) for 90 min (pulse), washed and then left for 1 h (chase). Cells were then treated accordingly before live-imaging. To follow cathepsin activity and lysosomal movement, 1 h before live-imaging the cells, Magic Red (Bio-Rad, Hercules, CA, USA) was added following manufacturer’s recommendation. Cells were then imaged for 2 min at a 2 s image frequency. To assess lysosomal membrane rupture, L6 cells were transfected to overexpress Galectin-3-GFP. Cells were then treated with iron or with LLoMe (1 h) prior to fixation with PFA and imaging.

### GST Recombinant Protein Expression and Purification

GST-empty vectors (ampicillin-resistant) and GST-RILP-C33 (truncated mutant) were expressed in BL21 competent cells and cultured overnight at 37 °C with shaking at 200 rpm. Once the OD reached 0.6, 1 µM isopropyl-β-D-thiogalactoside (IPTG) was added, and the cultures were incubated for 4 hours at 30 °C with shaking at 200 rpm. The induced recombinant GST-RILP-C33 and GST proteins were centrifuged at 5,000 ×g for 30 min at 4 °C. The resulting pellets were resuspended in bacterial lysis buffer (50 mM Tris, 150 mM NaCl, 10 mM β-mercaptoethanol, pH 7.5) supplemented with a protease inhibitor cocktail (Sigma-Aldrich). The bacterial lysates were then homogenized using an ultrasonic homogenizer (25% amplitude, with 30-second pulse on/off cycles) and centrifuged at 5,000 ×g for 30 min at 4 °C. The supernatants were incubated with a 50% glutathione-Sepharose 4B bead suspension prepared according to the manufacturer’s instructions (Millipore Sigma, G317-0756-01), at a dilution of 1:50 (beads to bacterial lysate supernatant, respectively), for 2 hours at 4 °C with gentle shaking. The samples were subsequently centrifuged at 5,000 ×g for 5 min at 4 °C, and the resulting pellets were resuspended in bacterial lysis buffer. The samples were analyzed by Western blotting to confirm protein expression and purification.

### Affinity Precipitation of GTP-Bound Rab7

L6 cells were treated and lysed in RIPA buffer (25 mM Tris-HCl, 150 mM NaCl, 1% Triton X-100, 1% sodium deoxycholate, pH 7.4) supplemented with a protease inhibitor cocktail (Sigma-Aldrich). The cells were incubated on ice for 30 min, collected into clean microcentrifuge tubes, and centrifuged at 3,000 ×g for 5 min at 4 °C. Before centrifugation, 10% of each lysate was reserved as an input control. The input samples were also centrifuged at 3,000 ×g for 5 min at 4 °C and subsequently mixed with an equal volume of 2× Laemmli sample buffer (0.5 M Tris, pH 6.8, glycerol, 10% sodium dodecyl sulfate (SDS), and β-mercaptoethanol at a 1:10 ratio). Previously prepared recombinant GST-RILP-C33 and GST proteins bound to glutathione-Sepharose 4B beads were mixed with the cell lysates at a dilution of 1:14 (beads to cell lysate, respectively). The mixtures were incubated for 2 hours at 4 °C with gentle shaking. The samples were then centrifuged at 500 ×g for 5 min at 4 °C and eluted using RIPA buffer at a 1:2 ratio (sample to buffer, respectively). The supernatants from each wash were transferred to clean microcentrifuge tubes, mixed with an equal volume of 2× Laemmli sample buffer, boiled at 95 °C for 5 min, and resolved by SDS-PAGE.

### Cell lysates, SDS-PAGE, and Western Blotting

L6 cells were lysed in RIPA lysis buffer (25 mM Tris pH 7.5, 150 mM NaCl, 1% Triton X-100, 1% Na Deoxycholate, 1% Protease Inhibitor). After denaturing lysates in Laemmli sample buffer for 10 min at 90 °C, samples were resolved on an SDS-PAGE gel and transferred onto polyvinylidene difluoride membrane (PVDF). The membranes were blocked with 3% bovine serum albumin (BSA) for 1 h at room temperature. Membranes were then incubated overnight at 4 °C in primary antibodies. Lysates were immunoblotted with the following primary antibodies from Cell Signaling Technology: Opa1 (1:1000, D6U6N rabbit monoclonal), Mfn2 (1:500, D2D10 rabbit monoclonal), p-Drp1 S616 (1:1000, rabbit polyclonal, CST #3455), total Drp1 (1:1000, D6C7 rabbit monoclonal), β-actin-HRP conjugate (1:1000, D6A8 Rabbit Monoclonal), GAPDH (1:1000, 14C10 rabbit monoclonal), and VDAC (1:1000, D73D12 rabbit monoclonal), Rab7 (1:2500, D95F2 rabbit monoclonal, cat #93675), β-actin (1:2500, rabbit polyclonal, cat #4967). Mfn1 (1:1000, EPR21953-74 rabbit monoclonal) was purchased from Abcam. Cathepsin D (1:1000, D-7 mouse monoclonal) was purchased from Santa Cruz Biotechnology (Dallas, TX, USA). GST (1:2500, rabbit polyclonal, cat A190-122A) was purchased from Bethyl Laboratories. Membranes were then washed and incubated in the corresponding horse-radish peroxidase (HRP)-linked secondary antibody from Cell Signaling Technology for 1 h at room temperature: anti-mouse IgG, HRP-linked antibody (1:10,000, horse polyclonal) and anti-rabbit IgG, HRP-linked antibody (1:10,000, goat polyclonal). The antibody-antigen complex was reacted with enhanced chemiluminescence (Bio-Rad) reagents before developing. Semi-quantitative densitometric analysis was performed using ImageJ software (NIH, Bethesda, MD, USA).

### Cell viability and Lactate Dehydrogenase (LDH) release assays

Cell viability was quantified using the ReadyProbes™ Cell Viability Imaging Kit (Thermo Fisher Scientific) according to the manufacturer’s protocol. Quantification was expressed as the percentage of non-viable cells (green fluorescence) relative to the total number of cells (blue fluorescence). Lactate dehydrogenase (LDH) release was quantified using the CytoScan™-LDH Cytotoxicity Assay Kit (G-Biosciences, Overland, MO, USA) in accordance with the manufacturer’s instructions.

### Immunofluorescence

Cells were fixed in 4% paraformaldehyde (PFA) in PBS for 10 min and then quenched with 100 mM glycine. After 3 washes with PBS, cells were permeabilized using 0.5% triton X-100 (Sigma-Aldrich) in PBS for 5 min, washed 3 times with PBS and blocked for 1 h with 1% BSA in PBS. Cells were then incubated with the following primary antibodies for 1 h: anti-TOMM20 (1:500, ST04-72 rabbit monoclonal, Invitrogen Cat.# MA5-32148), anti-TFEB antibodies (1:300, rabbit polyclonal, Bethyl cat A303-673A) or anti-Rab7 (1:200, D95F2 rabbit monoclonal, cat #93675). This was then washed 3 times with PBS for 5 min and incubated with the appropriate secondary antibodies for 1 hour: anti-rabbit DyLight-488 (1:500, Bethyl, donkey polyclonal). Then, cells were washed 3 times in PBS, labelled with 1 µg/mL DAPI (Roche) for 5 min, washed again with PBS and rinsed with distilled water. Finally, coverslips were mounted on slides with DAKO mounting media (Agilent, Santa Clara, CA, USA), and fluorescence images were obtained as described above.

### Fluorescence microscopy

Three microscopy systems were used in this study. One was an Olympus IX83 P2ZF spinning-disk confocal microscope system (Quorum Technologies, Inc., Puslinch, ON, Canada) equipped with a Hamamatsu camera (ORCA-Flash4.0 V3 Digital CMOS Camera). Images were acquired using a 60X oil immersion objective (UPLAPO OHR, 1.5 NA), and using filter sets according to the fluorescent dyes and proteins used: SD-DAPI (ex. 405; em. 440/40), SD-GFP (ex. 470; em. 510/50), SD-RFP (ex. 555; em. 600/50), SD-Cy5 (ex. 640; em. 700/75), SD-mt-Keima-green (ex. 470; em. 600/50). For live-cell imaging, coverslips were enclosed in a Leiden chamber and mounted in a microscope-mounted environmental chamber maintained at 37 °C and 5% CO_2_. The second system was a Quorum Diskovery spinning disc confocal microscope equipped with a Leica DMi8 microscope connected to an Andor Zyla 4.2 Megapixel sCMOS or an iXON 897 EMCCD camera and controlled by Quorum Wave FX powered by MetaMorph software (Quorum Technologies, Inc.). The third system was a Nikon A1 confocal microscope built on a Nikon Eclipse Ti2 inverted microscope stand (Nikon Instruments Inc., Melville, NY, USA), equipped with an environmental incubation chamber (37 °C and 5% CO_2_). Image acquisition and analysis were performed using NIS-Elements software (Nikon Instruments Inc.)

### Image Analysis

To assess mitochondrial fission in 2D, mitochondrial branch length and networking parameters were quantified as in (Valente et al., 2017). For 3D structural analysis, z-stacks were captured with a step size of 0.2 µm. The images were rendered using the surfaces tool on IMARIS software (Oxford Instruments, Abingdon, UK) to evaluate mitochondrial sphericity, where a value approaching 1 indicates a perfect spherical shape.

For L6::mt-Keima cells, to extract mitolysosomes using FIJI/ImageJ, the exact same protocol was followed as published by Kelly *et al*. (Kelly et al., 2024). Briefly, a mask was created using only the “red” excitation image with the “MaxEntropy Threshold” setting and then used to remove the background. Next, using the “Image Calculator” function, the “red” image was subtracted from the “green” image. The result was binarized, and the number of points was evaluated using the “Analyzed Particles” function. The number of dots per cell represent what was called “mitolysosome”.

To count the number of mCherry-Parkin puncta, FIJI/ImageJ was used. For each cell individually, “Analyze Particle” was run to count the number of particles with a size of 8 pixel^2^ to infinity, excluding edges. The distribution of mCherry-Parkin within cells was quantified as follows: cells displaying less than 10 Parkin puncta with a mean cytosolic intensity above 800 A.U. (background) were considered as “cytosolic only”; cells with more than 10 puncta and a cytosolic intensity below 800 A.U. (background) were considered as “puncta only”; cells with more than 10 puncta and a cytosolic intensity above 800 A.U. (background) were considered as “mixed cytosolic and punctate”.

### Quantitative Real-Time PCR (qPCR)

Total RNA was isolated using the RNeasy Mini Kit (Qiagen, Toronto, ON, Canada) and subsequently reverse-transcribed into cDNA utilizing the RevertAid RT Kit (Thermo Fisher Scientific). Real-time qPCR tracking was executed with SsoAdvanced Universal SYBR Green Supermix (Bio-Rad) using a Bio-Rad CFX384 Touch Real-Time PCR Detection System. The amplification program consisted of an initial activation step at 95 °C for 15 min, followed by 40 cycles of denaturation at 94 °C for 15 s, annealing at 55 °C for 30 s, and extension at 70 °C for 30 s. Relative transcript abundance was calculated using the comparative ΔΔCT method and normalized to beta-actin expression levels. The rat-specific primer sequences utilized were as follows: CtsB (Cathepsin B): Forward (5’-3’) TGGGTTCAGCGAGGACAT; Reverse (5’-3’) ATGGTGTAGGGTAAGCAGCC. CtsD (Cathepsin D): Forward (5’-3’) GCGTCTTGCTGCTCATTCT; Reverse (5’-3’) AACTCTGACACTGGCTCCTT. Beta Actin: Forward (5’-3’) AAGATCCTGACCGAGCGTGG; Reverse (5’-3’) CAGCACTGTGTTGGCATAGAGG.

### Statistical analysis

Statistical analyses were performed using GraphPad Prism following its recommendations for statistical tests. Data are presented as mean ± standard deviation or mean ± SEM, as indicated in the figure legends. The number of independent biological replicates, cells analyzed per condition, and statistical tests used for each experiment are reported in the corresponding figure legends. For microscopy-based experiments, cell measurements of a treatment group within each independent experiment were averaged to generate a single sample-level mean. The sample means from each independent experiment were then used for statistical comparison. When data are assumed to be normally distributed, we used parametric tests: paired t-tests were used for comparisons between two groups, while one-way repeated measures ANOVA were used for comparisons among three or more groups, and two-way ANOVA when two independent variables were compared, followed by the recommended post-hoc tests as specified in the figure legends. If data were normalized, then non-parametric tests were employed. Differences were considered statistically significant at p < 0.05, but p values are disclosed.

## Acknowledgements

We thank Dr. Ella Hyatt for managing the Toronto Metropolitan University-MaRS Research Facilities. This work was supported by funds from the Canada Research Chair program to RJB (950-232333), the Canadian Institutes of Health Research to RJB (PJT-183914), the Canada Foundation for Innovation to RJB (38151), contributions from Toronto Metropolitan University and Faculty of Science to RJB, and a Natural Sciences and Engineering Research Council of Canada to GW. The authors did not use generative AI tools to draft and write the manuscript. Google’s Gemini was used at the final draft stage to identify potential language errors like spelling or grammatical errors.

## Supplemental Information

**Supplemental Figure 1. ML098 promotes Rab7 membrane association. A.** Representative confocal microscope images of L6 cells treated or not with ML098 and then labelled with anti-Rab7 antibodies. **B.** Quantification of the number of Rab7 puncta. Data are shown as mean ± SEM from 3 independent experiments where at least 30 cells per condition per experiment were quantified. Data were tested using paired Student’s t-test.

