## Supplementary figures and images for "Rab7 activation ameliorates impaired lysosome dynamics and mitophagic flux during chronic iron overload"

### Supplemental Figure S1

**Sup. Fig 1. - ML098 is activating Rab7**

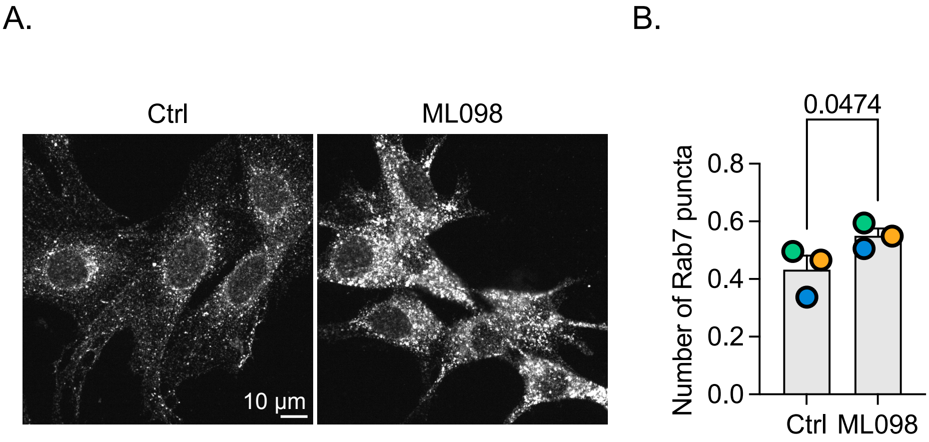
